# Evolution of chlorhexidine susceptibility and of the EfrEF operon among *Enterococcus faecalis* from diverse environments, clones and time spans

**DOI:** 10.1101/2022.01.27.478125

**Authors:** Ana P. Pereira, Patrícia Antunes, Rob Willems, Jukka Corander, Teresa M. Coque, Luísa Peixe, Ana R. Freitas, Carla Novais

## Abstract

Chlorhexidine (CHX) is widely used to control the spread of pathogens (e.g. human/animal clinical settings, ambulatory care, food industry). *E. faecalis*, a major nosocomial pathogen, is broadly distributed in diverse hosts and environments facilitating its exposure to CHX over the years. Nevertheless, CHX activity against *E. faecalis* is understudied. Our goal was to assess CHX activity and the variability of ChlR-EfrEF proteins (associated with CHX tolerance) among 673 field isolates and 1784 *E. faecalis* genomes from PATRIC database from different sources, time spans, clonal lineages and antibiotic resistance profiles. CHX minimum inhibitory concentrations (MIC_CHX_) and minimum bactericidal concentrations (MBC_CHX_) against *E. faecalis* presented normal distributions (0.5-64 mg/L). However, more CHX tolerant isolates were detected in the food chain and recent human infections, suggesting an adaptability of *E. faecalis* populations in settings where CHX is heavily used. Heterogeneity in ChlR-EfrEF sequences was identified, with isolates harboring incomplete ChlR-EfrEF proteins, particularly the EfrE identified in the ST40 clonal lineage, showing low MIC_CHX_ (≤1mg/L). Distinct ST40-*E. faecalis* subpopulations carrying truncated and non-truncated EfrE were detected, the former being predominant in human isolates. This study provides a new insight about CHX susceptibility and ChlR-EfrEF variability within diverse *E. faecalis* populations. The MIC_CHX_/MBC_CHX_ of more tolerant *E. faecalis* (MIC_CHX_=8mg/L; MBC_CHX_=64mg/L) remain lower than in-use concentrations of CHX (>500mg/L). However, CHX increasing use combined with concentration gradients occurring in diverse environments potentially selecting multidrug-resistant strains with different CHX susceptibilities, alert to the importance of monitoring the trends of *E. faecalis* CHX tolerance within a One-Health approach.

## INTRODUCTION

Chlorhexidine (CHX) is a disinfectant and antiseptic used since the 1950s and included in the World Health Organization’s list of essential medicines (1, 2). It has been widely used for different purposes (e.g. surface disinfectants, antiseptics, mouthwashes, personal care products) in hospitals, the community, food industry, animal husbandry and pets (3). Currently, CHX is recommended in the prevention of skin or oral colonization and consequent health care-associated infections by multidrug-resistant (MDR) bacteria, such as methicillin-resistant *Staphylococcus aureus* (MRSA) and vancomycin-resistant *Enterococcus* (VRE) (4, 5). As a bisbiguanide, CHX interacts with the cell wall and membrane anionic sites affecting the osmotic equilibrium of the cell, resulting in a bacteriostatic or bactericidal action depending on the concentration applied (2, 3, 6). Recommended CHX concentrations in disinfectants and antiseptics are usually high (0,05% and 4%; 500 to 40,000 mg/L) (2). However, CHX’s wide use has also negative effects, including ecotoxicity to aquatic life, horizontal transfer promotion of genetic elements carrying antimicrobial resistance genes, or changes in bacterial communities (e.g. in the oral microbiota towards a greater abundance of Firmicutes) (7–9).

Within Firmicutes, *Enterococcus* spp. is one of the most frequently found taxa in both humans and animals (10). They are members of the oral and gut microbiota of mammals, birds and reptiles, able to cause infections in animals, and one of the leading causes of human hospital-acquired infections globally (10). Their ability to tolerate different stresses facilitates their survival in the environment, being frequently recovered from plants and vegetables, water bodies and soil (10, 11). Also, this ability to survive and persist in abiotic surfaces is of particular concern in hospitals, increasing the risk of their transmission to patients followed by potential colonization or infection (12).

*Enterococcus faecium* populations of clade A1, a cluster overrepresented by clinical isolates, have shown a trend towards CHX tolerance (13). Strains belonging to this clade carry a single amino acid change (P102H) in a conserved DNA-binding response regulator (ChtR) from the 2CS-CHX^T^ operon (13, 14). CHX tolerance in *Enterococcus faecalis* remains, however, scarcely explored. Most available studies are restricted to clinical isolates, especially causing oral infections, and do not analyze the clonal diversity of the studied isolates (15–17). Recently, the *efrEF* operon, coding for the heterodimeric ATP-binding cassette (ABC) transporter EfrEF, was shown to be involved in the tolerance of the *E. faecalis* V583 strain to CHX (18). The EfrEF transporter is composed by the EfrE and EfrF proteins, and their upregulation under CHX exposure is mediated by ChlR, a putative MerR family transcription regulator (18, 19).

Our aim was to evaluate CHX susceptibility, the variability of the *chlR-efrEF* genes and to link CHX phenotypes with *chlR-efrEF* genotypes among a large collection of *E. faecalis* isolates from human, animal, food and environmental sources and available genomes from the last century. CHX activity results will be also discussed within the *E. faecalis* population structure context.

## RESULTS

### Chlorhexidine susceptibility of *E. faecalis* from diverse sources and clonal lineages

The minimum inhibitory concentration(s) of CHX digluconate (MIC_CHX_) of the 151 *E. faecalis* ranged from 0.5 to 8 mg/L, with an MIC_50_ of 4 mg/L and MIC_90_ of 8 mg/L (Fig. 1A). The highest MIC_CHX_ of 8 mg/L was observed in 21% (n=32/151) of the population studied while 6% (n=9/151) of isolates showed an MIC_CHX_ of 0.5-1 mg/L, corresponding in both cases to *E. faecalis* recovered from different sources and clonal lineages. MIC_CHX_ values presented a normal distribution, with a selected log_2_ standard deviation (SD) of 0.52 and a fitted curve overlapping the raw count distribution (Fig. 1A). The MIC ECOFF_CHX_ proposed for 99% of the population by the ECOFFinder tool was 8 mg/L. However, the MIC_CHX_ distribution analysis using the NORM.DIST Excel function showed a 4% probability of a wild-type isolate having an MIC_CHX_ of >8 and ≤16 mg/L and 0% >16 mg/L. Therefore, based on the normal distribution data, a tentative MIC ECOFF_CHX_ of ≤16 mg/L is suggested for *E. faecalis*.

**Fig 1.**
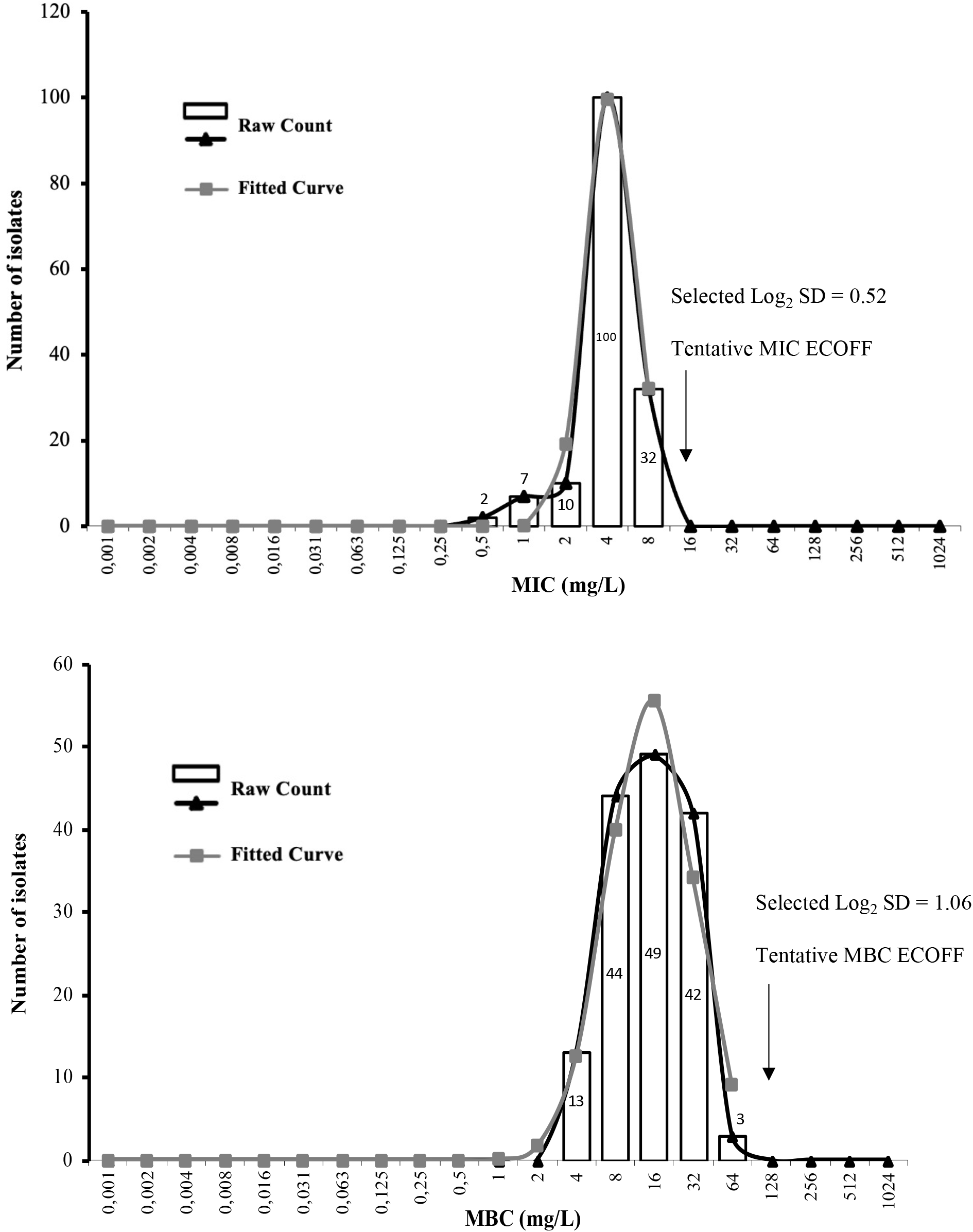
Distribution of the *Enterococcus faecalis* studied by different chlorhexidine minimum inhibitory concentrations-MIC (A) and minimum bactericidal concentrations-MBC (B). The graph fitted curves were estimated using the ECOFFinder tool which proposed 8 and 64 mg/L for MIC and MBC, respectively, as limits of 99% of wild-type population. The NORM.DIST Excel 16.44 function indicates that the probability of occurrence of an isolate with an MIC>8 and ≤16mg/L is 4% and 0% >16mg/L and with an MBC≤64mg/L is 100% and 0% >64mg/L. The tentative ECOFFs for MIC and MBC suggested are, therefore, 16 and 64 mg/L, respectively.

CHX digluconate minimum bactericidal concentration(s) (MBC_CHX_) ranged from 4 to 64 mg/L, with an MBC_50_ of 16 mg/L and MBC_90_ of 32 mg/L. A normal MBC_CHX_ distribution was also observed, being the selected log_2_ SD of 1.06 (Fig. 1B). The highest MBC_CHX_ of 32-64 mg/L (30%; n=45/151) as well as the lowest MBC_CHX_ of 4-8 mg/L (38%; n=57/151) comprised in both cases isolates from different sources and clonal lineages. The MBC ECOFF_CHX_ proposed for 99% of the population by the ECOFFinder tool was 64 mg/L, and the NORM.DIST Excel function estimated an 12% probability of a wild-type isolate having an MBC_CHX_ =64 mg/L and 0% >64 mg/L. Thus, both analysis point to a tentative MBC ECOFF_CHX_ of ≤64 mg/L for *E. faecalis*.

The analysis of CHX activity regarding isolates’ antibiotic resistance profiles showed that MDR *E. faecalis* had higher mean MIC_CHX_ but similar mean MBC_CHX_ comparing to non-MDR ones [5.0 vs 4.2 (*P*≤0.05) and 16.1 vs 19.4 mg/L (*P*≥0.05), respectively]. The MIC_CHX_ and MBC_CHX_ among VRE was variable and ranged, respectively, between 4-8mg/L and 4-32 mg/L (n=14; human infection, hospital sewage, human faecal samples at hospital admission and dog faeces from 1996-2016). MIC_CHX_/MBC_CHX_ of linezolid-resistant isolates varied between 1-8 mg/L and 16-64 mg/L (n=6; raw frozen pet food in 2019-2020), respectively.

### *E. faecalis* isolates from the food chain and recent human samples express higher tolerance to chlorhexidine

The MIC_CHX_ and MBC_CHX_ distribution of the 151 *E. faecalis* isolates tested were analyzed separately by source and time span (5-year intervals). The MIC_CHX_ distribution of the 151 *E. faecalis* revealed that the mean MIC_CHX_ of isolates from humans (4.8 mg/L; 44 STs among 77 isolates) was higher than the associated with isolates from the food chain (4.1 mg/L; 47 STs among 59 isolates) (*P*≤0.05) but similar to those from the environment (4.8 mg/L; 11 STs among 12 isolates) (*P*≥0.05). Within the group of *E. faecalis* from humans, the mean MIC_CHX_ was significantly higher among those associated with infection (5.4 mg/L; 27 STs among 41 isolates) than colonization (4.2 mg/L; 29 STs among 36 isolates) (*P*≤0.05). In contrast, mean MBC_CHX_ values were significantly higher among isolates from the food chain (22.6 mg/L) than isolates from humans or the environment (15.3 mg/L and 13.0 mg/L, respectively) (*P*≤0.001). MBC_CHX_ of *E. faecalis* from human infection or colonization isolates were similar (17.1 mg/L vs 13.2 mg/L, respectively; *P*≥0.05).

Food chain *E. faecalis* from different time spans showed variable MIC_CHX_ and MBC_CHX_, with no apparent increasing trend over time (Fig. 2A). However, a significant increasing trend in the mean MIC_CHX_ and MBC_CHX_ over the years was detected in isolates from human sources (Fig. 2B) (*P*≤0.05). We also analyzed the MIC_CHX_/MBC_CHX_ trends separately for strains associated with human infection or colonization (including isolates mostly from faeces or the urinary tract of healthy humans, but also faeces from long-term care facility patients and individuals at hospital admission) (Table S1). The mean MIC_CHX_ and MBC_CHX_ of isolates obtained from human colonization in 2001-2005 (3.8 and 10.8 mg/L, respectively; 15 STs among 16 isolates) was statistically similar to that of more recent ones (2016-2020: 4.2 and 16.8 mg/L; 13 STs among 16 isolates) (*P*≥0.05), although an increase was observed (Fig. 2C). In isolates from human infections the mean MIC_CHX_/MBC_CHX_ significantly increased, with the mean MBC_CHX_ tripling between 2001-2005 (10.5 mg/L; 12 STs among 13 isolates) and 2016-2020 (32.0 mg/L; 10 STs among 11 isolates) (*P*≤0.05) (Fig. 2D).

**Fig 2.**
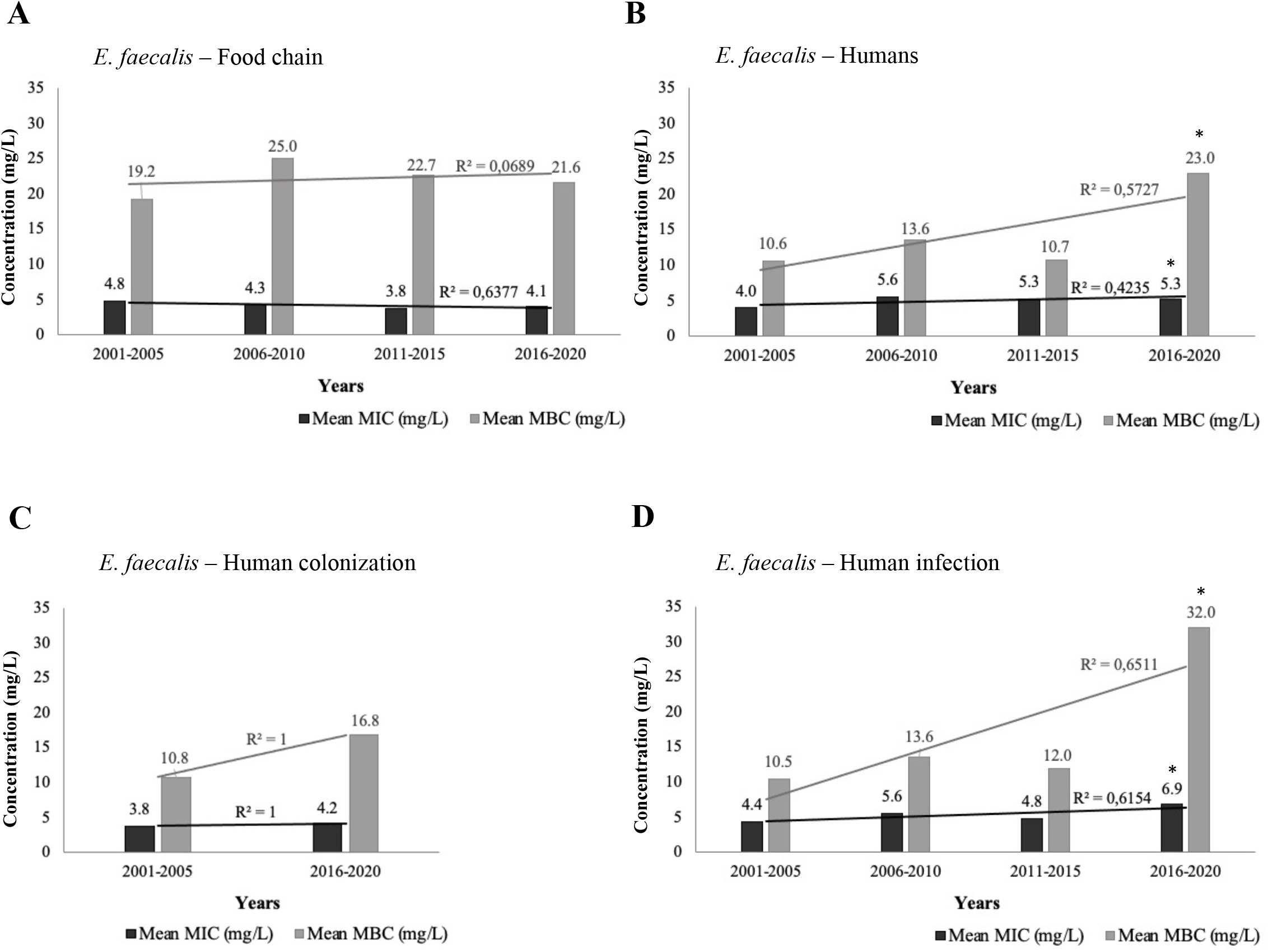
Chlorhexidine mean minimum inhibitory concentrations (MIC) and minimum bactericidal concentrations (MBC) distribution over the years (five-year intervals, from 2001 to 2020) of *E. faecalis* from independently analysed sources. (A) Distribution of food chain *E. faecalis* isolates (n=57). (B) Distribution of *E. faecalis* isolates recovered from all human sources (n=75). (C) Distribution of *E. faecalis* isolated from human colonization (including isolates from healthy-humans, long-term care patients and human faecal samples at hospital admission) between 2001-2005 and 2016-2020 (n=32). (D) Distribution of *E. faecalis* from human infection (n=39). *, *P*≤0.05; two-tailed unpaired Student’s *t*-test. *E. faecalis* from earlier years, between 1996 and 2000 (n=4), human colonization isolates from 2006-2015 (n=4) and those with other origins (n=15) were not included in the analysis due to the low number of isolates. A linear trendline and the R-squared (R^2^) value were added to each distribution using Excel 16.44.

### Diversity of ChlR-EfrEF sequences and association of incomplete proteins with *E. faecalis* low MIC_CHX_ values

The *efrEF* operon was identified in all but one of the 666 *E. faecalis* genomes analyzed, with 5% (n=33/666) carrying genes coding for incomplete ChlR (n=2), EfrE (n=25) or EfrF (n=6) proteins (Fig. S1 and 3, Table S2). In order to better recognize the impact of the incomplete ChlR, EfrE and EfrF proteins on the susceptibility to CHX, the MIC_CHX_ and MBC_CHX_ were also determined for all isolates with incomplete proteins which were not included in the group of 151 isolates formerly tested in the MIC_CHX_/MBC_CHX_ assays. Whereas the MIC_CHX_ values of most of these strains were consistently low (0.5-1 mg/L for 91% of the strains, n=30/33), the MBC_CHX_ values ranged from 1 mg/L to 64 mg/L, similarly to that observed for other isolates without frameshift, non-frameshift or nonsense mutations in the ChlR-EfrEF proteins (Table S2, Fig. 3).

**Fig 3.**
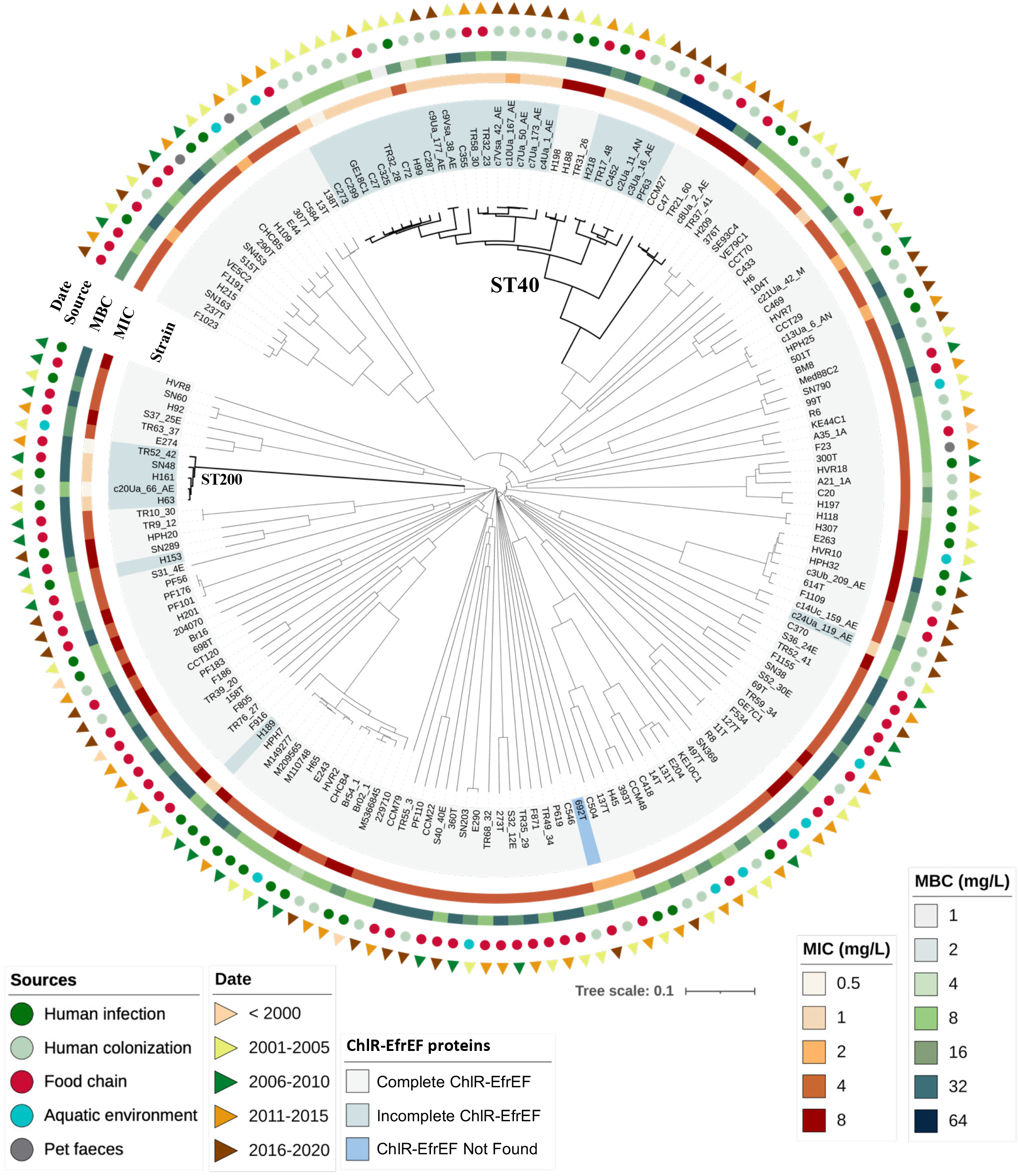
Phylogenetic tree based on the core genome MLST (cgMLST) allelic profiles of all sequenced *Enterococcus faecalis* studied with phenotypic assays (n=174). The clonal relationship of the strains was established from the sequence analysis of 1,972 gene targets accordingly to the *E. faecalis* cgMLST scheme (40), using Ridom SeqSphere+ software version 7.2. *E. faecalis* isolates’ features, marked with different colours and shapes using the iTol software (https://itol.embl.de), from the inner to the external part of the phylogenetic tree are: complete, incomplete or not found ChlR-EfrEF proteins marked in the “Strain” line, chlorhexidine minimum inhibitory concentrations (MIC), chlorhexidine minimum bactericidal concentrations (MBC), the source and date of isolation. For more isolates’ details see Table S2.

Among the 33 *E. faecalis* with incomplete ChlR-EfrEF, 25 isolates carrying a truncated EfrE and recovered from different sources belonged to ST40 (Table S2, Fig. 3). All of them showed a missing guanine in the nucleotide position 186 of the *efrE* gene associated with a frameshift mutation resulting in a stop codon at amino acid 79 of EfrE (Fig. S1, Tables S2 and S3). The search for common mutations in the PATRIC database available genomes showed that 85% (n=76/89) of the published ST40 *E. faecalis* also carried this *efrE* mutation (Fig. 4, Table S3). Proteins 100% identical to the truncated EfrE of ST40 *E. faecalis* were also found in 5 ST268 *E. faecalis* human faecal isolates (GenBank acc. numbers: NZ_CABGJG000000000; CABGJA000000000; BJTJ00000000; BJTS00000000; BJTH00000000).

**Fig 4.**
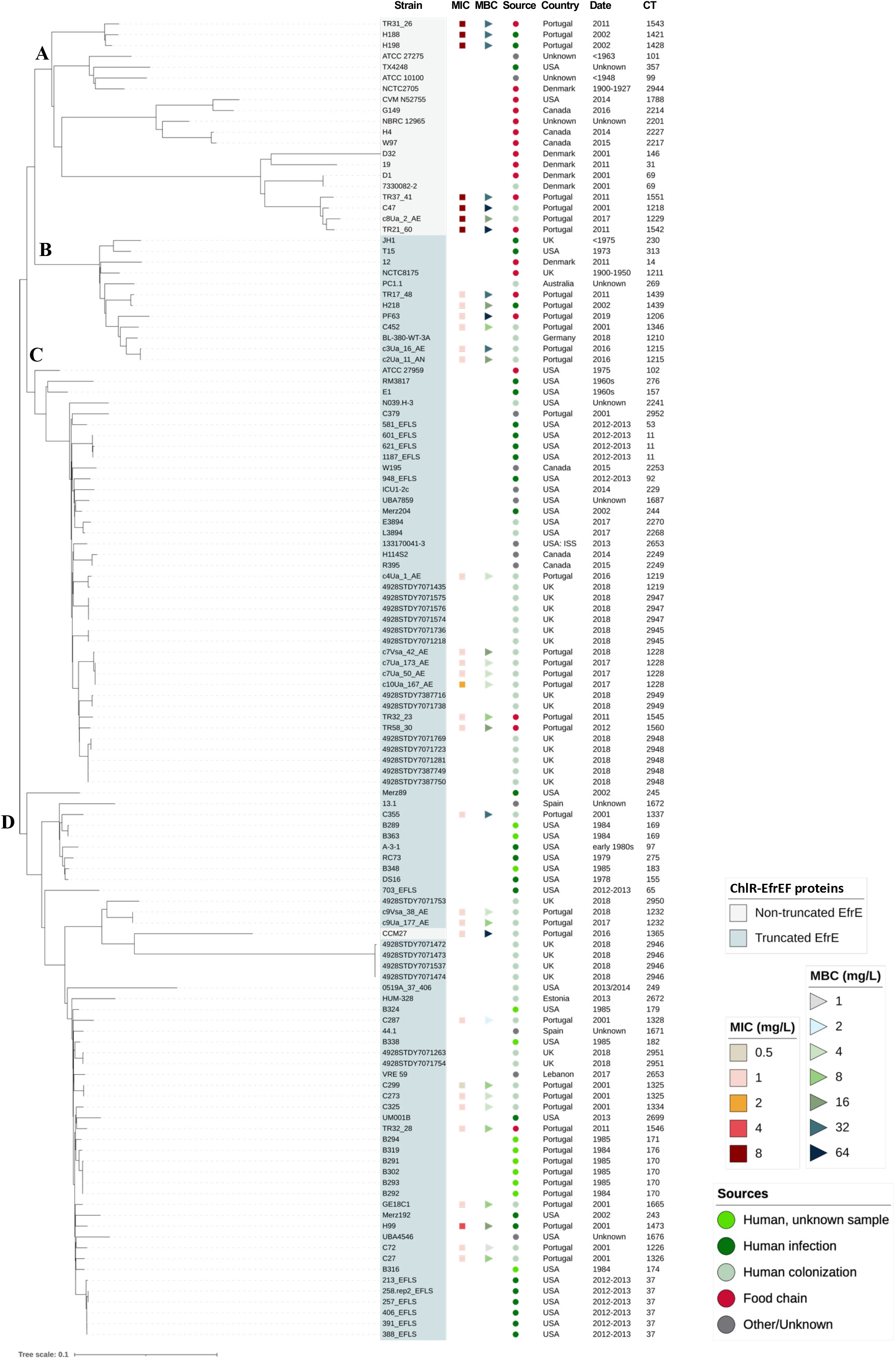
Phylogenetic tree based on the core genome MLST (cgMLST) allelic profiles of *Enterococcus faecalis* identified as ST40 from our collection and available at the PATRIC database (until the 18^th^ of December 2020) (n=122). The clonal relationship of the strains was established from the sequence analysis of 1,972 gene targets accordingly to the *E. faecalis* cgMLST scheme (40), using Ridom SeqSphere+ software version 7.2. Four clusters (A, B, C and D) were identified. Strains with a truncated EfrE (marked in the “Strain” column), chlorhexidine minimum inhibitory concentrations (MIC) and minimum bactericidal concentrations (MBC), the source and date of isolation as well as the CT (Complex Type) of each isolate were marked with different colours, shapes or by text, using the iTol software (https://itol.embl.de). For more isolates’ details see Table S3.

*E. faecalis* with incomplete ChlR or EfrF proteins were less represented in our collection (Table S2) as well as in the *E. faecalis* genomes searched in the PATRIC database. Concerning ChlR mutations, two human isolates from our collection (ST59 and ST319) showed the deletion of an adenine in *chlR* nucleotide 5 associated with frameshift mutations resulting in an early truncated protein at amino acid 7 (Fig. S1, Table S2). One published ST40 *E. faecalis* (food chain), with the previously described truncated EfrE, also showed an incomplete ChlR protein due to the insertion of an adenine in *chlR* nucleotide 530, resulting in an early truncated protein at amino acid 181 (Table S3).

Concerning EfrF mutations, an isolate from our collection presented a nonsense mutation in *efrF* (C1567T) resulting in an early stop codon at amino acid 523 in a single ST179 fecal isolate. This mutation was not found in other 30 ST179 human isolates analyzed (15 from our collection and 15 from PATRIC database) (Fig. S1, Tables S2 and S3). In addition, a deletion of 39 nt (696-734 nt) resulting in a shortened EfrF protein without amino acids from positions 233 to 245 was found in all ST200 analyzed (5 from our collection and 1 available at PATRIC database; 3 human and 3 food chain isolates) (Fig. S1). Finally, one public ST40 *E. faecalis* from human origin with the truncated EfrE protein had an EfrF with a frameshift mutation, caused by the insertion of an adenine in *efrF* nucleotide 1138, associated with an early stop codon at 392 amino acid position (Table S3).

Among the 632 isolates with a complete ChlR-EfrEF proteins, a broad range of missense mutations was identified in each of the proteins studied, but no correlation between specific mutations and MIC_CHX_ and/or MBC_CHX_ was noted (Table S2).

### EfrE-truncated ST40 *E. faecalis* clustered separately from non-truncated ST40 ones in the phylogenetic tree and were mostly recovered from humans

To establish an association between clonal lineages and CHX phenotypes, we performed the core-genome multi-locus sequence typing (cgMLST) based phylogeny of all sequenced *E. faecalis* isolates with available phenotypic information (n=174). We identified 77 STs and 160 complex types (CTs) with variable MIC_CHX_ and MBC_CHX_ values for isolates of each ST or CT (Fig. 3; Table S2). Nonetheless, it is of note that ST40 *E. faecalis* (18 CTs) and ST200 *E. faecalis* (5 CTs) isolates expressing lower MIC_CHX_ (0.5-1 mg/L) clustered separately, while the few ST179, ST308 and ST319 with low MIC_CHX_ were dispersed throughout the phylogenetic tree (Fig. 3).

To further analyze the ST40 *E. faecalis*, all 33 ST40 genomes from our collection and the 89 available at the PATRIC database (n=122) were separately analyzed in a new cgMLST-based phylogenetic tree (Fig. 4). Isolates with operons encoding a truncated EfrE protein clustered separately from those with operons encoding a complete EfrE protein. Cluster “A” grouped 20 of the 21 strains with a non-truncated EfrE (Fig. 4, Table S3), whereas ST40 *E. faecalis* with a truncated EfrE grouped in clusters “B” (n=12 isolates), “C” (n= 39 isolates) or “D” (n=50 isolates), the latter comprising also one isolate with non-truncated EfrE. The oldest *E. faecalis* with a truncated EfrE was recovered from the food chain in 1900-1950. Overall, ST40 *E. faecalis* with a truncated EfrE included in clusters C and D were isolated predominantly from humans (81%; n=82/101; *P*<0.0001) of different geographical regions. ST40 *E. faecalis* isolates from cluster A had an MIC_CHX_ of 8 mg/L while most ST40 isolates of clusters B, C and D (n=24/26) had an MIC_CHX_ of 1 mg/L. The only ST40 *E. faecalis* with non-truncated EfrE included in cluster D presented the same ChlR-EfrEF mutations as a ST308 *E. faecalis* from a healthy human, which also had an MIC_CHX_ of 1 mg/L without possessing an incomplete ChlR-EfrEF (Table S2; Fig. 3). Additionally, most of our isolates of clusters A and B had an MBC_CHX_ ≥16 mg/L (92.3%, n=12/13; *P*<0.0001), while strains in clusters C and D mostly had an MBC_CHX_ <16 mg/L (75%, n=15/20; *P*≤0.05).

## DISCUSSION

The increasing challenge to control the growth and transmission of human and animal pathogens in clinical settings, ambulatory care or in the food industry explains the rising use of biocides in different sectors, namely of CHX. However, the scarcity of available data concerning both wild-type bacterial phenotypes and subpopulations’ adaptation to biocides over the years limits the perception and the restraint of a potential biocide resistance threat.

In this study we showed that the MIC_CHX_ and MBC_CHX_ normal distributions for the *E. faecalis* isolates analyzed were in accordance with the ranges previously reported for this species (15, 20). However, the higher mean MBC_CHX_ values found in isolates from the food chain as well as the increasing mean MIC_CHX_/MBC_CHX_ values of recent isolates from human infections suggest the adaptability of *E. faecalis* populations in settings where CHX is heavily used. A tentative MIC ECOFF_CHX_ and MBC ECOFF_CHX_ of 16 mg/L and 64 mg/L, respectively, proposed by the ECOFFinder tool and the NORM.DIST Excel function analysis based on *E. faecalis* normal distribution, seems, therefore, limited because it comprises isolates with heterogeneous phenotypes and genotypes. Although further molecular analysis are needed to understand the significance of such diversity in bacterial populations classified as “wild-type” for CHX, the MIC/MBC_CHX_ values found are considerably below the in-use concentrations of CHX (500 to 40,000 mg/L) (2, 3). Nevertheless, they are within or higher than the levels that have been detected in the skin of patients subjected to CHX bathing (<4.69-600 mg/L), in cow milk (4-78 mg/L) or in sewage (28-1300 ng/L) (5, 21, 22). As CHX tends to persist in water, sediment and soils (23), diverse *E. faecalis* populations showing different CHX susceptibilities could hypothetically be selected and adapt within gradients of sub-inhibitory concentrations occurring not only in patients’ skin but also in diverse environments (5, 21–25).

The detection of *E. faecalis* isolates falling into the upper borderline of the MBC_CHX_ distribution (32-64 mg/L), with many of them recently recovered from human infections or the food chain, and some showing resistance to vancomycin or linezolid, alerts for the possibility of MDR strains selection by CHX as well as an adaptation towards CHX tolerance in the following years. Such increase in CHX tolerance over time has been described for other relevant bacterial species, such as *Staphylococcus aureus, Klebsiella pneumoniae* or *Acinetobacter baumanni* (26–28), suggesting that the increasing use of CHX since the 2000s in community, veterinary and hospital contexts (22, 27) might have been contributing to this ecological adaptation. Moreover, other bacterial stresses, as those with impact in membrane fluidity (e.g. temperature, acids, other biocides) should also be considered in future studies to assess cross-tolerance with CHX (29, 30), and to help explain the higher MBC_CHX_ found in isolates from the food chain throughout the study, when comparing to isolates from humans sources, more tolerant to CHX in recent years.

The few articles addressing the genetic mechanisms involved in CHX tolerance among *E. faecalis* described the upregulation of different genes, especially the conserved *chlR-efrEF* genes (18). We showed that *chlR-efrEF* diversity does not seem to have a direct impact in the MBC_CHX_ values, but variants with incomplete proteins encoded by *chlR-efrEF* affect *E. faecalis* growth at low CHX concentrations (corresponding to MIC_CHX_), particularly in ST40 *E. faecalis* from humans. ST40 *E. faecalis* are known to be widely distributed in different environments and hosts (31), but a divergent evolution among strains with truncated and non-truncated EfrE was detected, being both selected across different time spans and geographical regions. Most *E. faecalis* with truncated EfrE, presenting the same mutation, were of human origin, being isolated from this source at least since the 1960’s. However, whether this truncated EfrE subpopulation reflects multiple evolved genomic regions of ST40 *E. faecalis* with a better human host adaptation, namely to colonization, remains to clarify. More studies are also needed to better understand the role of the EfrEF operon in the metabolism of *E. faecalis* and specifically in the tolerance to CHX and other stresses, as this operon was described to be involved in the transport of ethoxylated fatty amines, fluoroquinolones and fluorescent dyes (18, 19, 32). Although changes in the *chlR-efrEF* genes seem to impact strains’ growth inhibition by CHX in most cases, a few isolates (ST40, ST59 and ST860) with incomplete/deleted ChlR-EfrEF exhibited MIC_CHX_ levels >1 mg/L suggesting the occurrence of other cellular mechanisms allowing bacteria growth under CHX exposure.

In conclusion, our study provides novel and comprehensive insights about CHX susceptibility within the *E. faecalis* population structure context, revealing more CHX tolerant subpopulations recovered from the food chain and recent human infections. Although a functional EfrEF operon was previously described to be important to *E. faecalis* V583 response to CHX (18), we further show a detailed analysis of the genetic diversity of the operon and the correlation with CHX phenotypes, namely the impact of incomplete ChlR-EfrE proteins on isolates’ growth (MIC_CHX_). The recent strains with a higher tolerance to CHX and the known multiple sources for CHX diffusion pollution (e.g., down-the-drain of CHX containing products used in diverse society sectors) (23) alert for the potential consequences of the growing CHX use and to the need of continuous monitoring *E. faecalis* adaptation towards CHX tolerance within a One Health approach.

## METHODS

### Epidemiological background of field isolates included in the different assays

A collection of 673 *E. faecalis* isolates (666 sequenced), representative of different geographical regions, sources, time spans, and genomic backgrounds (BioProjects PRJEB28327; PRJEB40976 and PRJNA663240) (31, 33), was selected for this study. They were recovered in previous studies from human infection (n=174), human colonization (n=163), food chain (animal production settings, animal meat and other food products) (n=275), pets (n=9) and aquatic environment (n=45) samples, in diverse regions (Portugal, Tunisia, Angola, Brazil) and time spans (1996-2020) (33–35). Among them, 181 isolates were included in the CHX susceptibility assays (details in Table S1), with 41% (n=75/181) classified as MDR (resistance to 3 or more antibiotics from different families), 8% (n=14/181) as resistant to vancomycin and 3% (n=6/181) to linezolid, in previous studies (33–35). Of these, 151 *E. faecalis* were initially considered to evaluate *E. faecalis* MIC_CHX_/MBC_CHX_ distributions. Subsequently, 30 additional *E. faecalis* with ChlR-EfrEF incomplete proteins and/or belonging to ST40 were considered for phenotypic-genotypic comparative studies along with the former 151 isolates. These 30 additional strains were not included in the first set of phenotypic assays not to introduce an overrepresentation of *E. faecalis* with ChlR-EfrEF incomplete proteins and/or belonging to ST40 in MIC_CHX_/MBC_CHX_ distributions.

### Chlorhexidine susceptibility

The MIC_CHX_ (CAS: 18472-51-0, Sigma Aldrich) of the 181 *E. faecalis* was established by broth microdilution, using the methodological approach proposed by the Clinical and Laboratory Standards Institute (CLSI) for antimicrobial susceptibility testing (Muller-Hinton broth; pH 7.4; 37°C/20h) (36). Using a 96-well microtiter plate containing serial two-fold dilutions of CHX (concentration range of 0.125 to 128 mg/L), bacterial suspensions in logphase growth, adjusted to reach a final inoculum of 5×10^5^ CFU/ml in each well, were incubated for 20h at 37°C. Microdilution panels were prepared before each assay. The first concentration of CHX without visible growth was considered the MIC_CHX_. Pinpoint growth was often observed and disregarded as recommended (36).

To determine the MBC_CHX_, 10μl of each well without visible growth from the 96-well MIC_CHX_ plate were incubated onto brain heart infusion (BHI) agar plates at 37°C for 24h, as defined by the CLSI (37). The MBC_CHX_ was defined as the lowest CHX concentration for which the number of colonies was equal or less than the rejection value defined by CLSI guidelines, based on the final inoculum of each well confirmed by actual count (37). Each experiment was repeated 3-6 times and the MIC_CHX_/MBC_CHX_ values corresponded to the mean of the determinations. *E. faecalis* ATCC 29212 and *E. faecalis* V583 strains were used as controls.

The assessment of MIC_CHX_ and MBC_CHX_ wild-type distribution was performed using the ECOFFinder tool (ECOFFinder_XL_2010_V2.1; available at http://www.eucast.org/mic_distributions_and_ecoffs/), which attempts to fit a log-normal distribution to the presumptive wild-type counts by the so-called iterative statistical method (38). In order to increase specificity to identify wild-type strains, the percentage selected to set the ECOFF was 99%, as suggested by the guidelines of the ECOFFinder tool. The NORM.DIST Excel v.16.44 function was used to calculate the probability of occurrence of isolates at higher concentrations and, consequently, evaluate the potential presence of an acquired tolerance mechanism if such probability was too low, using the mean, the standard deviation and with the cumulative normal distribution function option set to TRUE (38). Finally, the statistical significance of the differences between MIC_CHX_ and MBC_CHX_ of isolates from the diverse sources, time spans and with disparate antibiotic resistance profiles was assessed using the two-tailed unpaired Student’s *t*-test (Excel v.16.44), and the differences associated with the source and MBC_CHX_ distribution among *E. faecalis* ST40 populations were analyzed by the Fisher exact test using GraphPad Prism software, version 9.0., with *P* values ≤0.05 considered significant.

### Whole-genome sequence analysis

The genomic search of *chlR*, *efrE* and *efrF* genes (reference strain: *E. faecalis* V583; GenBank accession no. AE016830.1; locus-tag EF_2225 to EF_2227) was performed in the 666 *E. faecalis* sequenced genomes by using the MyDBfinder tool available at the Center for Genomic Epidemiology (www.genomicepidemiology.org). The *chlR-efrEF* genes identified in each genome were translated into the corresponding amino acid sequences by the DNA translate tool of ExPASy SIB Bioinformatics Resource Portal (https://web.expasy.org/translate/) and the occurrence of incomplete ChlR-EfrEF proteins was evaluated.

For the sequenced *E. faecalis* included in the phenotypic assays, a comparison of the amino acid sequences with the reference strain *E. faecalis* V583 was performed using the Clustal Omega software (https://www.ebi.ac.uk/Tools/msa/clustalo/) to identify specific mutations. Their clonal relationship was also established by MLST and cgMLST ((39); http://pubmlst.org; (40); Ridom SeqSphere+, version 7.2). A phylogenetic tree based on their cgMLST allelic profiles was constructed using Ridom SeqSphere+ software and isolates’ information was added to the tree using the iTol software (https://itol.embl.de).

### Comparative genomics

In order to evaluate the frequency of strains with genes coding for incomplete ChlR, EfrE or EfrF proteins in other collections, ChlR, EfrE and EfrF sequences with 100% identity until the stop codon with those found in our isolates with incomplete ChlR-EfrEF were searched in 1784 *E. faecalis* genomes of the PATRIC database, representing a timespan between 1900-2020 (last update on the 18^th^ of December 2020). In addition, to assess if *E. faecalis* isolates containing genes encoding incomplete ChlR, EfrE or EfrF proteins had a similar genomic evolution, a cgMLST-based phylogenetic tree was constructed with all *E. faecalis* genomes identified as ST40 (n=122), both from our collection and available at the PATRIC database (last update on the 18^th^ of December 2020), using the Ridom SeqSphere+ software. Isolates’ information was added to the tree using the iTol software (https://itol.embl.de).

## Acknowledgments

This work is financed by national funds from FCT - Fundação para a Ciência e a Tecnologia, I.P., in the scope of the project UIDP/04378/2020 and UIDB/04378/2020 of the Research Unit on Applied Molecular Biosciences - UCIBIO and the project LA/P/0140/2020 of the Associate Laboratory Institute for Health and Bioeconomy - i4HB, by the AgriFood XXI I&D&I project (NORTE-01-0145-FEDER-000041) cofinanced by European Regional Development Fund (ERDF) and through the NORTE 2020 (Programa Operacional Regional do Norte 2014/2020). Ana Paula Pereira was supported by a PhD fellowship from FCT (SFRH/BD/144401/2019), co-financed by European Social Fund through Norte Portugal Regional Operational Programme (NORTE 2020); and Ana R. Freitas by the Junior Research Position (CEECIND/02268/2017 - Individual Call to Scientific Employment Stimulus 2017) granted by FCT/MCTES through national funds. The funders had no role in study design, data collection and interpretation, or the decision to submit the work for publication.

CRediT authorship contribution statement: Ana Paula Pereira: Methodology, Software, Formal analysis, Investigation, Writing – original draft, Writing – review & editing. Patrícia Antunes: Formal analysis, Investigation, Writing – review & editing, Funding acquisition. Rob Willems: Formal analysis, Writing – review & editing; Jukka Corander: Formal analysis, Writing – review & editing; Teresa M. Coque: Formal analysis, Writing – review & editing. Luísa Peixe: Supervision, Funding acquisition, Formal analysis, Writing – review & editing. Ana R. Freitas: Conceptualization, Methodology, Formal analysis, Supervision, Writing – review & editing, Funding acquisition. Carla Novais: Conceptualization, Methodology, Software, Formal analysis, Investigation, Supervision, Writing – original draft, Writing – review & editing, Funding acquisition, Project administration.

Authors have no conflicting interests relevant to the study.

